# TIRE-seq: an Integrated Sample Extraction and Transcriptomics Workflow for High Throughput Perturbation Studies

**DOI:** 10.1101/2024.11.20.624406

**Authors:** Piper O’Keeffe, Yasmin Nouri, Hui Shi Saw, Zachery Moore, Tracey M. Baldwin, Sam W.Z. Olechnowicz, Jafar S. Jabbari, David McG Squire, Stephen Leslie, Changqing Wang, Yupei You, Matthew E. Ritchie, Ryan S. Cross, Misty R. Jenkins, Cindy Audiger, Shalin H. Naik, James R. Whittle, Saskia Freytag, Sarah A. Best, Peter F. Hickey, Daniela Amann-Zalcenstein, Rory Bowden, Daniel V. Brown

## Abstract

RNA sequencing (RNA-seq) is widely used in biomedical research, advancing our understanding of gene expression across biological systems. Traditional methods require upstream RNA extraction from biological inputs, adding time and expense to workflows. We developed TIRE-seq (Turbocapture Integrated RNA Expression Sequencing) to address these challenges. TIRE-seq integrates mRNA purification directly into library preparation, eliminating the need for a separate extraction step. This streamlined approach reduces turnaround time, minimizes sample loss, and improves data quality. A comparative study with the widely used Prime-seq protocol demonstrates TIRE-seq’s superior sequencing efficiency with crude cell lysates as inputs. TIRE-seq’s utility was demonstrated across three biological applications. It captured transcriptional changes in stimulated human T cells, revealing activation-associated gene expression profiles. It also identified key genes driving murine dendritic cell differentiation, providing insights into lineage commitment. Lastly, TIRE-seq analyzed the dose-response and time-course effects of temozolomide on patient-derived neurospheres, identifying differentially expressed genes and enriched pathways linked to the drug’s mechanism of action. With its simplified workflow and high sequencing efficiency, TIRE-seq offers a cost-effective solution for large-scale gene expression studies across diverse biological systems.

## 1 Introduction

Transcriptomics has driven groundbreaking discoveries, playing a pivotal role in advancing our understanding of gene expression across diverse biological systems [1]. One major success of transcriptomics is its application in large-scale initiatives such as The Cancer Genome Atlas (TCGA) and Genotype-Tissue Expression (GTEx) project, providing deep insights into biology [2, 3]. Beyond consortium-scale efforts, transcriptomics has been employed in smaller studies to investigate processes underlying disease and development [4]. Despite its utility, the high cost of RNA sequencing (RNA-seq) presents a significant barrier to large-scale experiments. The expense of library preparation and sequencing limits the number of samples that can be analyzed, reducing the statistical power of these studies. Popular RNA-seq protocols, such as the Illumina TruSeq stranded RNA kit, require individual processing of each sample in separate reactions before applying unique indices during the final amplification step [5]. This workflow is both labor-intensive and resource-demanding, restricting scalability. Additionally these commercial kits lack flexibility, often requiring RNA extraction and normalization as prerequisites for library preparation.

Single-cell RNA-seq (scRNA-seq) methodologies have introduced innovations such as sample barcoding during reverse transcription, enabling early pooling [6]. Most scRNA-seq protocols sequence the 3’ or 5’ ends of transcripts, which reduces costs compared to full-length sequencing such as TruSeq. Incorporating unique molecular identifiers (UMIs) further enhances data quality by adjusting for PCR bias [7]. These innovations have inspired the development of several high-throughput, plate-based bulk RNA-seq protocols [8–16]. PLATE-seq was among the first bulk RNA-seq protocols to utilize early barcoding and sample pooling [8]. This method employs a Qiagen TurboCapture plate to purify mRNA from crude lysates in an automated format. The TurboCapture plate uses immobilized poly-T oligonucleotides to hybridize mRNA onto the solid surface of the plastic, followed by elution into a destination plate containing reverse transcription primers. Similarly, Prime-seq validated a magnetic bead-based RNA purification workflow upstream of library preparation steps and performed favorably compared to TruSeq [14]. PLATE-seq and Prime-seq both utilize 3’ transcript tag sequencing. 5’ sequencing, initially developed for transcription start site identification via cap analysis gene expression (CAGE) [17]—has seen less adoption. FACS-Based 5-Prime End Single-Cell RNA-seq (FB5P-seq) was developed for BCR/TCR repertoire analysis, placing sample barcodes and UMIs on a template-switching oligo (TSO) to accommodate early pooling [10]. Similarly, Survey of TRanscription Initiation at Promoter Elements with high-throughput sequencing (STRIPE-seq) employs barcoded TSOs for genome-wide transcription start site profiling [11].

Despite these advances, most RNA-seq protocols require RNA extraction as a distinct and time-consuming step prior to library preparation. This process demands careful handling to prevent RNA degradation and ensure data quality, posing challenges when processing large numbers of samples across multiple batches. An ideal bulk RNA-seq protocol would accommodate a wide range of input types, such as crude cell lysates and intact cells, enabling direct processing from cultured cells in microwell plates thus simplifying experimental workflows.

To address these limitations, we developed TurboCapture Integrated RNA Expression sequencing (TIRE-seq), a protocol that integrates RNA extraction directly with 5’ tag based library preparation. By eliminating the need for separate RNA purification, TIRE-seq streamlines workflows, reduces labor, and minimizes sample loss, offering a cost-effective and scalable solution for high-throughput transcriptomics.

## 2 Results

### 2.1 Integration of RNA Extraction and Library Preparation Using TIRE-seq

TIRE-seq leverages Qiagen’s TurboCapture plate to purify poly-A RNA directly from crude lysates, followed by reverse transcription and sample barcoding through immobilized poly-T oligos (Fig. 1a). Soluble single stranded DNA is then pooled and undergoes standard short-read based library preparation.

**Fig. 1.**
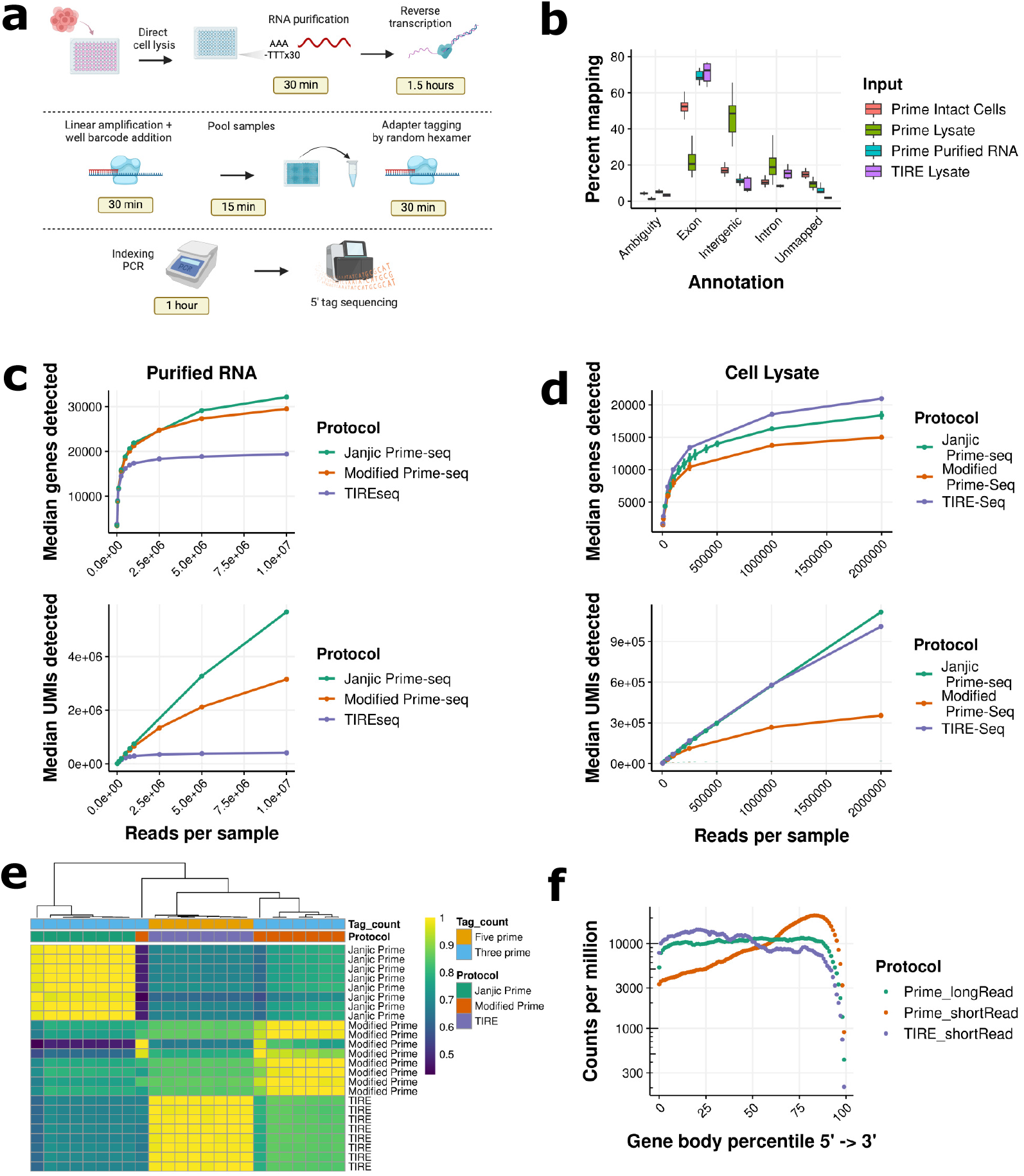
TIRE-seq outperforms Prime-seq on cell lysates. (a) Overview of the TIRE-seq workflow. Cells are lysed directly in media and transferred to Qiagen TurboCapture plates. mRNA hybridizes to immobilized poly-T oligos, followed by washing and reverse transcription. After well barcoding, samples are pooled, and library preparation occurs on the pooled material. Gene expression is quantified using 5’ tag sequencing. (b) Read mapping statistics for Prime-seq and TIRE-seq. “Intact cells” refers to FACS-sorted cells.”Lysate” refers to crude homogenized lysates, and “Purified RNA” refers to RNA purified using silica columns. (c) Downsampling analysis of 11 ng Universal Human Reference RNA (UHRR). Janjic = E-MTAB-10139. Top: Median number of genes detected. Bottom: Median number of unique molecules (UMIs) detected. Error bars represent the median absolute deviation of replicates. (d) Performance of TIRE-seq on 10,000 cell equivalents from HEK293T cell lysates. (e) Pearson correlation of logCPM for UHRR replicates across protocols. (g) Binned transcript coverage across protocols. ShortRead = Illumina 70-nt tag sequencing. LongRead = Oxford Nanopore sequencing of full-length Prime-seq cDNA.

We found the protocol to be sensitive at cell inputs ranging from 100,000 to 15,000 cells in a volume of 20 *μ*L (Fig. S1a and b). Despite being a pooled bulk RNA-seq protocol, sample cross-contamination was minimal (Fig. S1c). We assessed UMI saturation [18], finding that the highest-expressing gene in our datasets reached approximately 6,000 counts, well below the theoretical encoding capacity of 1,048,576 molecules for a 10-nt UMI (Fig. S1d).

Using crude lysates as input, TIRE-seq achieved higher exonic mapping compared to our modified implementation of Prime-seq. Interestingly, the TIRE-seq mapping rate with lysate as input was comparable to Prime-seq with purified RNA input (Fig. 1b).

We investigated the impact of lysis buffers and storage conditions on TIRE-seq data quality. Qiagen TCL buffer produced the highest-quality data, consistent with it being optimized for use with the TurboCapture plate (Fig. S1g). We stored cell lysates at 4 °C and - 80 °C for one week in lysis buffer. Superior data quality was observed with lysates stored at -80 °C, indicating sample degradation at 4 °C (Fig. S1e).

In a benchmarking experiment with Prime-seq, we used purified Universal Human Reference RNA (UHRR) as input (Fig. 1c). TIRE-seq exhibited poorer performance compared to our modified Prime-seq and the original Janjic dataset [14]. This was likely due the same RNA input amount in a 20 *μ*L input volume resulting in a 16-fold lower RNA concentration than Prime-seq. In contrast, with crude lysate input equivalent to 10,000 cells, TIRE-seq achieved a comparable number of detected molecules and higher gene detection compared to Janjic et al. (Fig. 1d). TIRE-seq also demonstrated greater molecular efficiency than our implementation of Prime-seq using the same sample. We compared cross-contamination rates between TIRE-seq and Prime-seq, observing lower contamination in TIRE-seq (Fig. S1h). TIRE-seq was highly reproducible across replicates and showed stronger correlation with our Prime-seq implementation than between the two Prime-seq protocols (Fig. 1e).

As a 5’ tag sequencing protocol, TIRE-seq exhibited broader transcript coverage with short-read sequencing, approaching the coverage achieved by Prime-seq when sequenced on an Oxford Nanopore instrument (Fig. 1f). This is likely to the presence of alternate transcriptional start sites [19].

### 2.2 Comparative Performance of TIRE-seq and Prime-seq in a High-throughput Experiment

As an initial biological application, human T cells were stimulated *in vitro* using CD3- and CD28-coated microbeads, followed by harvesting of CD4 and CD8 subsets at defined timepoints (Fig. 2a). For Prime-seq, RNA was extracted from cell lysates using magnetic beads, as described in *Methods*. In contrast, for TIRE-seq, cell lysates containing 4,000 cell equivalents were used directly without prior RNA purification.

**Fig. 2.**
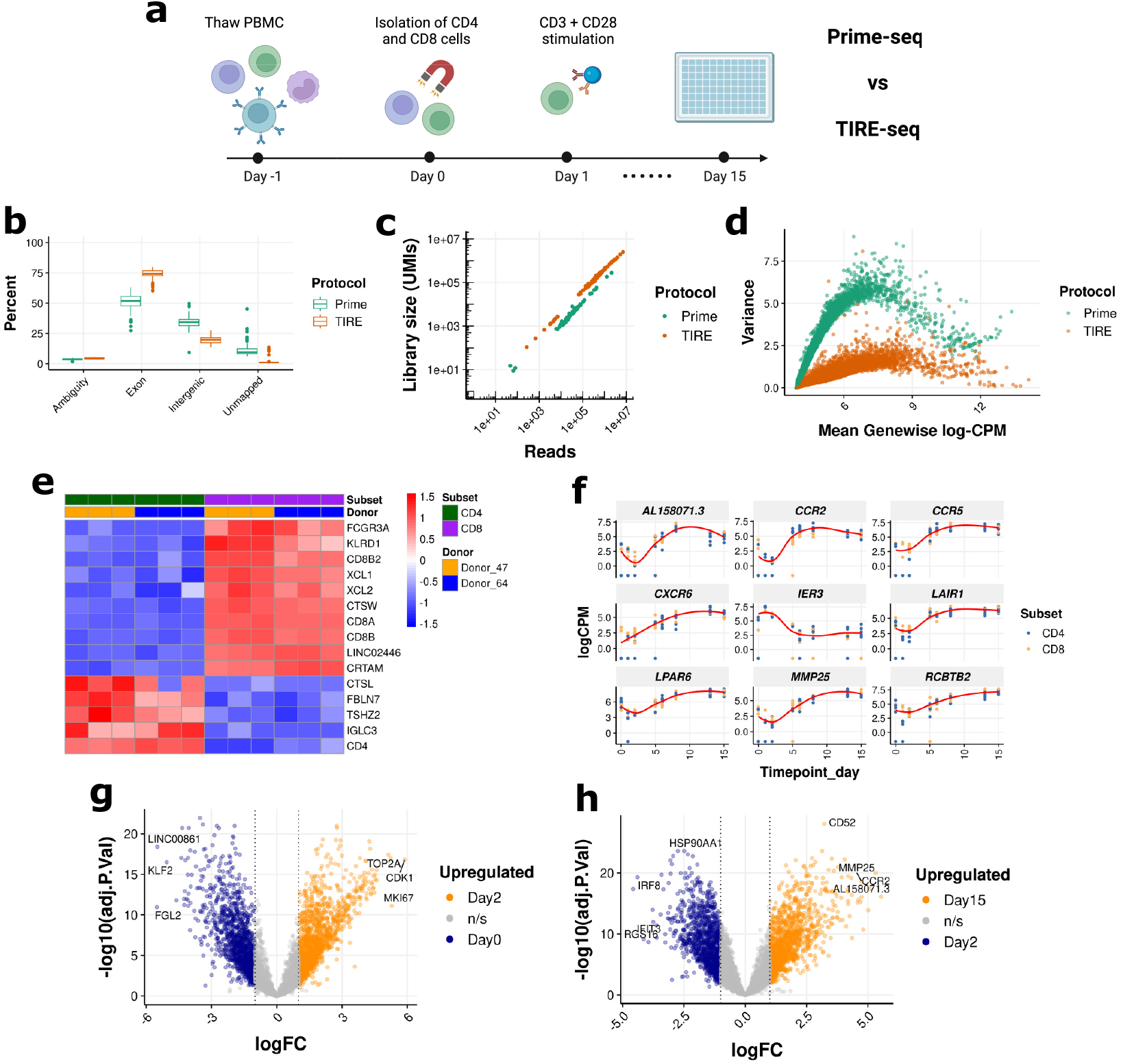
Superior performance of TIRE-seq compared to Prime-seq in a high-throughput experiment. (a) Experimental design of the human T cell experiment. T cell subsets were isolated on day 0, followed by stimulation on day 1. CD4 and CD8 subsets were sorted at the indicated timepoints, and both Prime-seq and TIRE-seq were performed on the same inputs. (b) Mapping rates to genomic features for Prime-seq and TIRE-seq. Reads were downsampled to the same value. (c) Molecular efficiency comparison between Prime-seq and TIRE-seq. Reads were downsampled to the same value. (d) Comparison of gene expression variability between Prime-seq and TIRE-seq. Reads were downsampled to the same value, with no gene filtering applied. (e) Heatmap showing logCPM values for the top 15 differentially expressed genes between CD4 and CD8 T cells at the day 2 timepoint. (f) logCPM values for the top six most variable genes over time during T cell stimulation. (g) Volcano plot of differentially expressed genes between day 2 and unstimulated (day 0) T cells in response to CD3 and CD28 stimulation. (h) Volcano plot of differentially expressed genes between day 15 and day 2 T cells in response to CD3 and CD28 stimulation.

Quality control metrics revealed superior exonic mapping for TIRE-seq compared to Prime-seq (Fig. 2b). TIRE-seq demonstrated enhanced sequencing efficiency by recovering a greater number of unique molecules at equivalent sequencing depth (Fig. 2c). Following stimulation, T cells exhibited a characteristic burst of transcriptional activation within 1–2 days [20]. This trend was observed in both protocols, with TIRE-seq displaying a greater number of counts at all timepoints (Fig. S2a), suggesting higher technical efficiency. The correlation between gene expression measurements from the two protocols was high, with a Pearson coefficient of 0.887 (Fig. S2b).

The improved technical metrics of TIRE-seq translated into reduced variability in gene expression measurements (Fig. 2d). Principal component analysis (PCA) of the samples revealed that early timepoints post-stimulation exhibited the most distinct transcriptional changes, while changes at later stages were less pronounced (Fig. S2c). Given the superior performance of TIRE-seq, these biological differences were more distinctly resolved in the PCA of TIRE-seq data, with a higher proportion of variance captured in the first two principal components (Fig. S2d). This enhancement led us to prioritize TIRE-seq data for downstream biological analyses. Differential expression analysis between CD4 and CD8 subsets at the peak of transcriptional activation on day 2 confirmed the expected expression patterns. CD8 subunits and CD4 markers were among the most differentially expressed genes (Fig. 2e). These findings align with the cells being sorted based on CD4 and CD8 surface marker expression (Fig. S2e).

The high-throughput nature of TIRE-seq enabled longitudinal profiling of gene expression, capturing dynamic responses to stimulation over a 15-day period (Fig. 2f). For genes such as the chemokine receptor *CCR2*, a transient decrease in expression during the activation phase was followed by a steady increase and subsequent plateau. This pattern reflects the dominance of a proliferative gene expression program at early timepoints (days 1–2) (Fig. 2g). This caused immune specific programs to appear diminished in relative terms (Fig. S2f).

In contrast, the Immediate Early Response 3 gene (*IER3*) was expressed at high levels at early timepoints, decreasing and stabilizing at day 7 (Fig. 2f). The elevated expression of *IER3* at day 0 prior to stimulation likely resulted from cellular stress induced by thawing from cryogenic storage. A comparative analysis between days 2 and 15 identified additional key regulators of immune responses, including Interferon Regulatory Factor 8 (*IRF8*) and Interferon-Induced Protein with Tetratricopeptide Repeats 3 (*IFIT3*), which showed reduced expression as the cells transitioned from peak activation to a steady-state (Fig. 2h). IFIT proteins are induced by interferon-*γ* and involved in the antiviral response [21]. Focusing on specific cytokines, the expression profiles of interleukin-2 (*IL-2*) and interferon-*γ* (*IFN-γ*) were consistent with expected activation kinetics: initial upregulation within 24 hours, followed by enhanced proliferative signaling at day 2 (Fig. S2f and g).

### 2.3 Gene Expression Dynamics in Murine Dendritic Cell Differentiation with TIRE-seq

Our next biological application of TIRE-seq employed a murine differentiation model. Bone marrow was harvested and hematopoietic stem cells were induced to differentiate into dendritic cells (DCs) using the ligand Flt3L (Fig. 3a). At the appropriate timepoints, DC subsets were sorted based on extracellular marker expression directly into 2x TCL buffer, bypassing the RNA extraction step that would be required with Prime-seq. TIRE-seq was performed on intact cell inputs consisting of 5,000 - 10,000 cells per well.

**Fig. 3.**
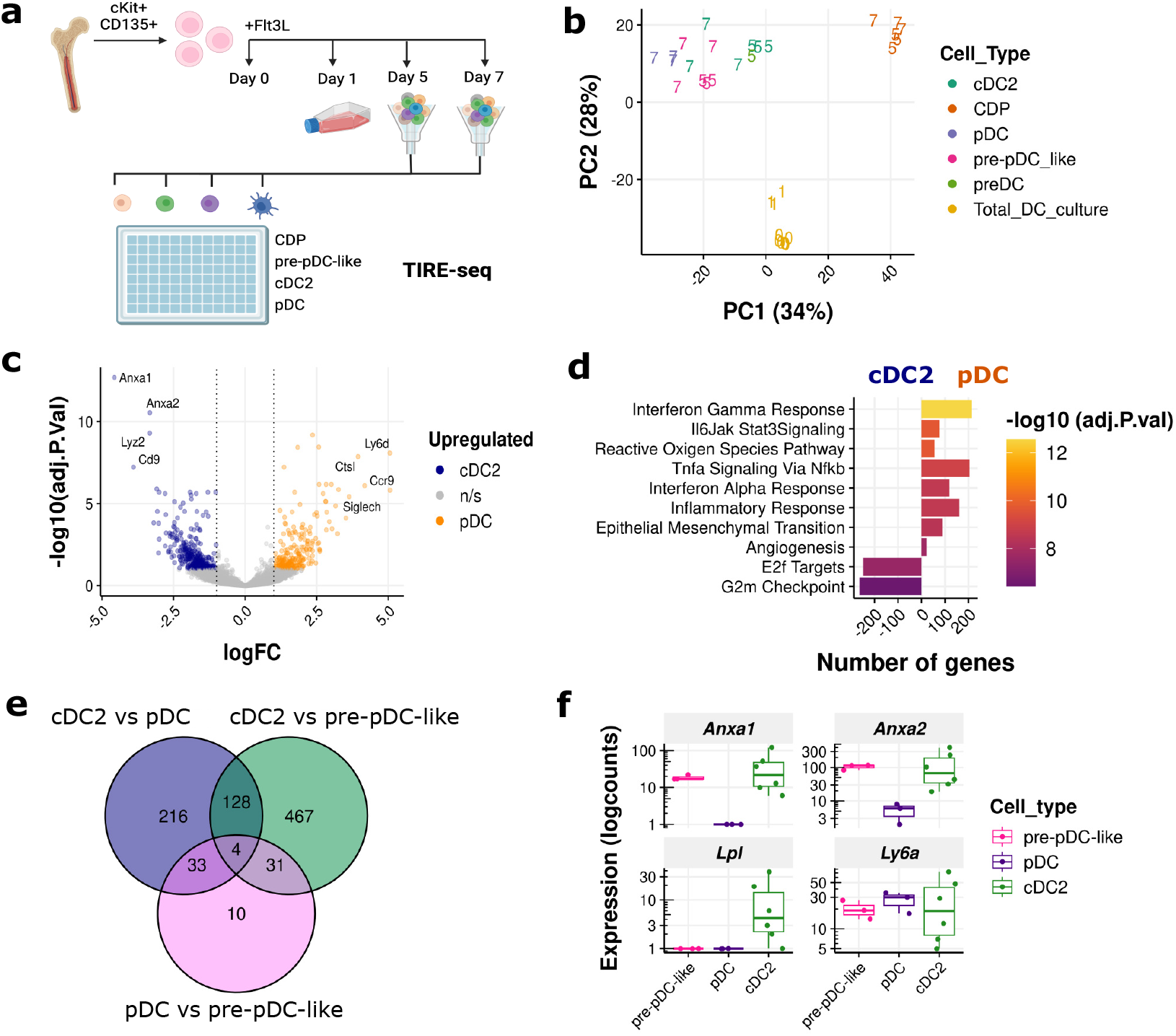
Comparative Transcriptomics of DC Subsets Using TIRE-seq. (a) Experimental design of the murine dendritic cell experiment. Hematopoietic stem cells were harvested from bone marrow and stimulated with Flt3L to induce dendritic cell differentiation. DC subsets were sorted into 96w plates for TIRE-seq at the indicated timepoints. (b) Principal component analysis (PCA) visualization of the dendritic cell experiment. The number of days post-Flt3L stimulation is annotated directly in the plot. (c) Volcano plot comparing cDC2 and pDC subsets seven days after Flt3L stimulation. (d) Gene set enrichment analysis of hallmark gene sets comparing pDCs and cDC2 cells. (e) Venn diagram showing the number of differentially expressed genes between the indicated DC subsets. (f) Expression of the four shared differentially expressed genes from panel e, visualized as box plots.

Samples clustered primarily according to their DC state, as determined by marker expression. The common dendritic progenitor (CDP) emerged as the most distinct population, consistent with its characterization as a poised cell state (Fig. 3b). While the post-Flt3L stimulation timepoint impacted some DC subsets, this effect was not uniform. In the differential expression analysis of two major DC lineages, type-2 conventional dendritic cells (cDC2) and plasmacytoid dendritic cells (pDCs), we identified established markers at the RNA level, including *Siglech* and *Ly6d* for pDCs, and *Cd9* for cDC2s (Fig. 3c). Ly6D surface marker staining confirmed these findings, whilst Siglec-H was one of the markers used in the flow cytometry sorting of pDCs (Fig. S3a).

In line with previous findings, inflammatory pathways were enriched in pDCs compared to cDC2s (Fig. 3d) [22]. A time-course analysis identified distinct dynamically regulated genes converging on antigen presentation pathways (Fig. S3b and c).

Of particular interest is the distinction between pre-pDC-like cells and more terminally differentiated pDCs and cDCs. There is evidence to suggest that pre-pDC-like cells can differentiate into cDC2s [23]. As expected, we found significantly more differentially expressed genes between cDC2s and pre-pDC-like cells than between pDCs and pre-pDC-like cells (Fig. 3e; Fig. S3d and e), Chi-square test *P <* 10^*−*14^.

Examination of the four genes that were consistently differentially expressed across these cell types revealed three genes exhibiting a gradient pattern of expression (Fig. 3f). Both Annexin A1 (*Anxa1*) and Annexin A2 (*Anxa2*) showed intermediate expression profiles in pre-pDC-like cells. Expression of these genes was elevated in cDC2s and reduced in pDCs. Annexin A1 plays a critical role in DC migration and the activation of inflammatory pathways [24].

In contrast, the elevated expression of *Ly6a*/Sca-1 aligns with previously reported increases during the differentiation of pre-pDC-like cells into pDCs [25].

### 2.4 TIRE-seq enables Temporal and Dose-Dependent Transcriptomic Analysis

We next applied a more complex experimental design, incorporating both a dose-response and time-course analysis within a 96-well plate (Fig. 4a). With this design it is convenient to use TIRE-seq by adding lysis buffer directly to each culture well at the appropriate timepoint. Patient-derived neurospheres (PDNs) are 3D cultures derived from resected tumor tissue and more accurately recapitulate glioblastoma multiforme (GBM) compared to traditional 2D cultures [26]. Temozolomide (TMZ) is the standard-of-care chemotherapeutic used clinically for patients with GBM [27].

**Fig. 4.**
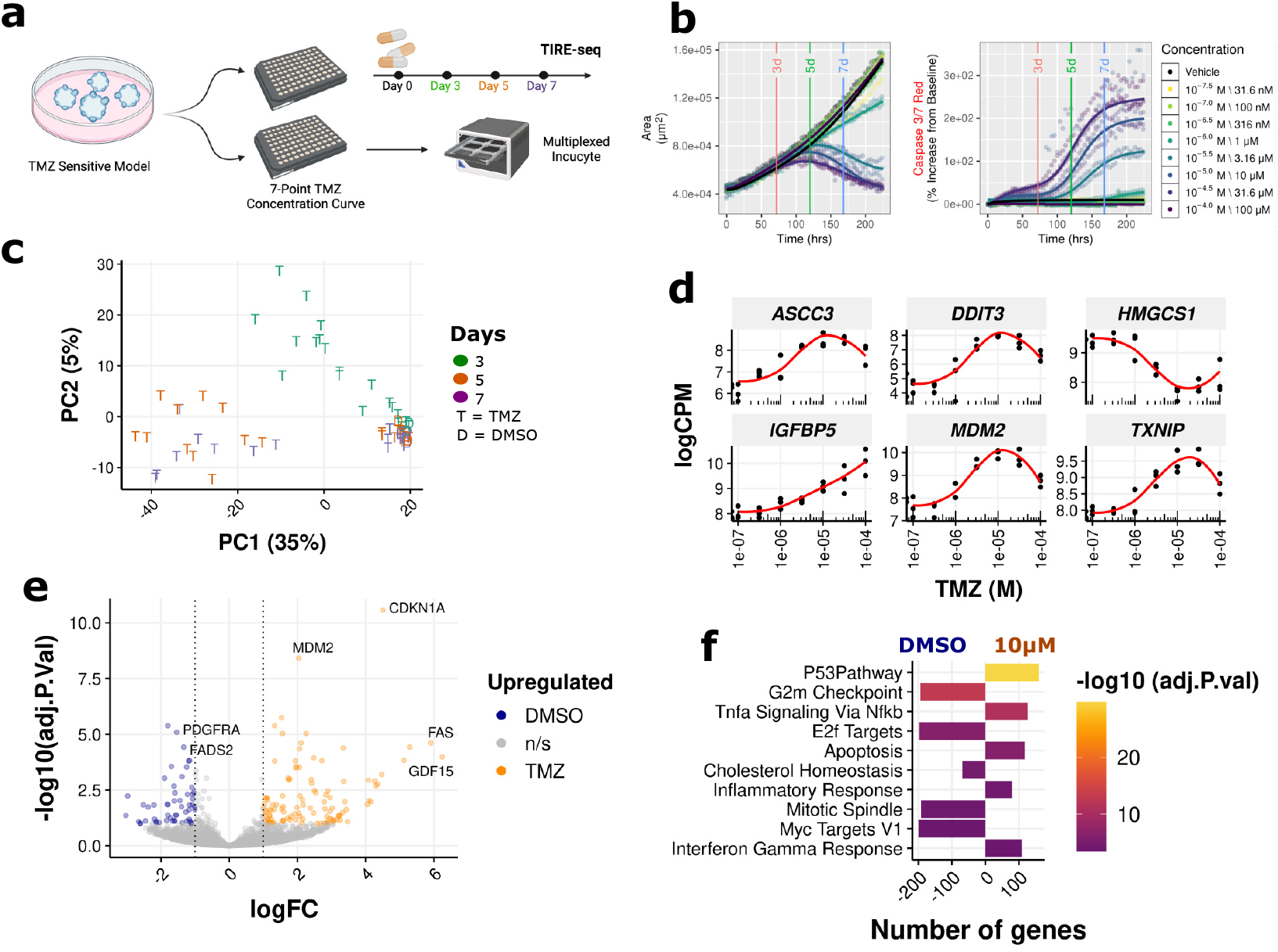
Tracking Temporal and Dose-Dependent Changes in patient-derived neurospheres with TIRE-seq. (a) Experimental design for TMZ-treatment of PDNs. Cells were exposed to varying doses of TMZ and sampled at three timepoints for TIRE-seq. A parallel culture was monitored for cell growth and death using IncuCyte imaging. (b) Growth curve (left) and caspase reactivity (right) from IncuCyte imaging. (c) Principal component analysis (PCA) visualization of the PDNs experiment. Samples are colored by timepoint and labeled by treatment: D = DMSO, T = Temozolomide. (d) LogCPM values of the top six most variable genes across timepoints across escalating TMZ doses. (e) Volcano plot of differentially expressed genes between 10 *μ*M TMZ and DMSO-treated neurospheres following three days of treatment. (f) Gene set enrichment analysis (GSEA) with hallmark gene sets comparing 10 *μ*M TMZ and DMSO-treated PDNs after three days of treatment.

During the time-course and dose-response experiment, PDNs were exposed to TMZ, while a parallel culture was monitored in real-time using the IncuCyte system to track proliferation and cell death (Fig. 4b). As PDNs proliferate slower than 2D cultures, no significant difference in growth was observed three days post-TMZ treatment. Minor growth differences began to emerge at 5 days, with clear distinctions observed at the 7 day endpoint. Coupled with the analysis of caspase-3 and caspase-7 cleavage activity as a proxy for cell death, we identified TMZ activity in this PDN model at concentrations above 3.16 *μ*M.

Principal component analysis revealed that TMZ treatment and timepoint were the dominant factors influencing gene expression (Fig. 4c), while the effect of TMZ dose was less pronounced (Fig. S4a). We next analyzed the genes that varied most with escalating TMZ doses after 3 days of treatment, before growth and viability effects became evident (Fig. 4d). The top dose-responsive genes were involved in cell stress and apoptosis, reflecting the switch from growth arrest to apoptosis at higher TMZ concentrations. Most genes showed a tapering off of expression at higher doses, potentially indicating a global shutdown of transcription.

Notably, the upregulated genes *ASCC3* and *DDIT3* are involved in DNA repair and endoplasmic reticulum stress respectively, highlighting pathways activated in response to TMZ’s genotoxic effects [28, 29]. Insulin-like Growth Factor Binding Protein 5 (*IGFBP5*) exhibited a near-linear increase in expression with escalating TMZ doses, consistent with its known role in promoting GBM stemness, survival, and invasion [30, 31]. Interestingly, while 3-hydroxy-3-methylglutaryl-CoA synthase 1 (*HMGCS1*) has been associated with a stem-like state in previous reports, it was downregulated in response to treatment [32].

Focusing on the 10 *μ*M dose at the three-day time-point, we found that the most upregulated genes were involved in cell cycle regulation and apoptosis, including *CDKN1A, FAS*, and *GDF15* (Fig. 4e). The most downregulated genes included *PDGFRA* and *FADS2*. Apoptotic and cell cycle arrest pathways were the most affected by TMZ treatment (Fig. 4f). Notably, cells exposed to TMZ at this early timepoint showed activation of TNF-*α* and Interferon-*γ* pathways. In a differential expression analysis, no significantly upregulated genes were observed when comparing cells treated with 100 *μ*M TMZ to those treated with 10 *μ*M (Fig. S4b). This finding suggests that while similar gene expression programs were active at both doses, they varied in magnitude (Fig. S4c and d).

We then performed a timepoint-centric analysis focusing on the 10 *μ*M dose (Fig. S4e). Due to sparse sampling across timepoints, we observed no prominent gene expression patterns. However, *CLU* (Clusterin) was identified as the most dynamically upregulated gene, while the chromatin-associated protein *HMGN2* was the most downregulated gene. Clusterin is a chaperone protein that has been observed to be upregulated in glioblastoma clinical samples [33]. When comparing the 3 day and 7 day timepoints, the most upregulated gene was *PBX1*, while the most downregulated gene was *CDCA5* (Fig. S4f). Upregulation of *PBX1* activates stem cell programs, while suppression of *CDCA5* induces cell cycle arrest [34, 35]. These findings suggest that cells surviving seven days of TMZ treatment may enter a quiescent, stem-like state [36].

## 3 Discussion

We developed a novel protocol, TIRE-seq, offering a streamlined RNA sequencing approach by integrating RNA extraction with library preparation. This workflow simplifies processing, reduces labor, and minimizes sample loss, making TIRE-seq a powerful tool for high-throughput transcriptomics. Previous studies have employed the Qiagen TurboCapture plate for purification, but mRNA was eluted into a separate plate for downstream processing [8]. Our innovation lies in combining both purification and library preparation within a single plate, further enhancing efficiency. TIRE-seq’s design significantly reduces turnaround time and achieves high sensitivity with low cell inputs, ranging from 100,000 to 4,000 cells in a 20 *μ*L volume. Although it operates as a pooled bulk RNA-seq protocol, sample cross-contamination remains minimal. This may be attributed to extensive washing steps, a specific hybridization sequence (template-switch oligo) instead of poly-T(30) and a shorter well-barcoding oligo, which are more effectively removed by SPRI bead-based size selection after pooling. Furthermore, TIRE-seq demonstrates superior sequencing efficiency compared to conventional protocols such as Prime-seq when using crude cell lysates as input (Table 1). In terms of transcript coverage, TIRE-seq outperforms Prime-seq with short-read sequencing and approaches the coverage obtained from Prime-seq using the Oxford Nanopore long-read platform. This advancement underscores TIRE-seq’s potential to transform large-scale transcriptomic studies by enabling faster, more cost-effective, and scalable analysis, making it an invaluable asset for both basic research and clinical applications.

**Table 1.** Summary of Experimental Workflow Differences: Prime-seq and TIRE-seq.

| Feature | Prime-seq | TIRE-seq |
| --- | --- | --- |
| Adherent cells | Washes required | Direct lysis |
| Suspension cells | Requires centrifugation | Direct lysis in suspension |
| Sample Volume | <1–5 $\mu L$ | <80 $\mu L$ |
| RNA Extraction | Separate upstream step | Integrated in workflow |
| Sequencing Efficiency | Lower | Higher |
| Genomic DNA contamination | Higher | Lower |
| Cost | Lower (ex. extraction) | Higher |
| Protocol Time | RNA extraction + 2 days | 1.5 days |
| Long Reads | Compatible | Incompatible |

In this study, TIRE-seq was applied to three biological contexts, demonstrating its capacity to capture dynamic transcriptional changes with high precision. In human T cells, TIRE-seq identified a robust transcriptional burst following stimulation, confirming its ability to capture gene expression dynamics. Differential expression analysis distinguished key genes separating CD4 and CD8 T cells, providing insights into their functional divergence. These results highlight the potential of TIRE-seq for immune profiling studies requiring dense temporal resolution to capture cell-specific responses. TIRE-seq also monitored gene expression during the differentiation of murine bone marrow cells into DCs. The data captured distinct expression profiles across DC subsets, validating known marker genes. Pathway analysis revealed a higher enrichment of inflammatory pathways in pDCs compared to cDC2s, consistent with the antiviral functions of pDCs. Time-course analysis further identified genes involved in antigen presentation, demonstrating TIRE-seq’s capability to track temporal changes throughout the differentiation process. Finally, TIRE-seq facilitated analysis of PDNs treated with TMZ. The protocol enabled simultaneous evaluation of dose and duration, revealing their significant impact on gene expression. Differential expression analysis identified genes involved in cell stress, apoptosis, DNA repair and stemness. These findings demonstrate the utility of TIRE-seq in drug response studies, especially in complex 3D cultures where precise temporal and dose-dependent gene expression patterns are critical for understanding therapeutic effects.

The strength of TIRE-seq lies in its compatibility with large volumes of cell lysates, typical of high-throughput cell culture experiments in multiwell plates. However, in a direct comparison with an equal quantity of purified RNA using Prime-seq, TIRE-seq exhibited poorer metrics. This is likely due to a 16-fold dilution of RNA relative to Prime-seq and the need for hybridization and reverse transcription on the solid surface of TurboCapture plates, as opposed to in-solution reactions in Prime-seq. We further discuss our benchmarking experiments in Supplementary Note D. Reagent volumes are higher for TIRE-seq, making it more expensive than Prime-seq and related methods, though still more cost-effective than TruSeq. We believe the reduction in hands-on time, achieved by integrating RNA extraction into the workflow, offsets the higher consumable costs, though this benefit may vary depending on experimental circumstances.

Since RNA purification and reverse transcription in TIRE-seq is performed using poly-A hybridization with prefabricated TurboCapture oligos, there is no universal method to introduce PCR handles on both ends of the cDNA strand. As a result, TIRE-seq is currently incompatible with long-read sequencing. Although the poly-T sequence can serve as a PCR primer binding site, this approach carries the risk of introducing internal priming artifacts [37]. Future development of TIRE-seq could focus on replacing TurboCapture plates with magnetic streptavidin poly-T beads would facilitate automation, reduce reagent volumes, and further lower costs. Additional savings could be achieved by developing gene-specific TIRE-seq using biotinylated gene-specific oligos within a streptavidin bead system.

TIRE-seq represents a significant advancement in high-throughput transcriptomics. Its streamlined workflow, high sensitivity, and improved data quality make it a valuable tool for moderate and large-scale studies.

## 4 Methods

All oligonucleotides were purchased at desalted grade from Integrated DNA Technologies unless otherwise specified. Sequences are provided in Table S1.

### 4.1 Ethical statement

PBMCs were isolated from unrelated healthy control donor samples obtained with consent from the Australian Red Cross lifeblood donor Registry (23-06VIC-16). The study was approved by WEHI Human Research Ethics Committee (17-01LR) and project 21/21. Mice (C57BL/6) were bred and maintained under specific pathogen-free conditions. Experiments were performed in accordance with institutional guidelines (WEHI Animal Ethics Committee 2021.051 and 2021.054).

### 4.2 Cell culture

HEK293T and NIH3T3 cells were grown and maintained in DMEM media (ThermoFisher A4192101) supplemented with 10% FBS, 5% sodium pyruvate and penicillin (100 I.U/mL) and streptomycin (100 *μ*g/mL). Jurkat and U937 cells were grown and maintained in RPMI media (ThermoFisher 11875093) supplemented with 10% FBS, 5% sodium pyruvate and penicillin (100 I.U/mL) and streptomycin (100 *μ*g/mL). Cells were maintained at 37 °C and 5% CO2. Cell lines were authenticated by STR typing (AGRF, Australia). Mycoplasma contamination was inspected with FastQ Screen analysis of the sequencing reads. All cells lines were negative. The BT504 PDN line used within this work was a kind gift from Keith Ligon, and was maintained as spheroids in NeuroCult NS-A Proliferation Media (STEMCELL Technologies 05751) supplemented with required additional growth factors (20 ng/mL human recombinant EGF, 10 ng/mL human recombinant bFGF, 2*μ*g/mL heparin, STEMCELL Technologies). Cultures were seeded on ultra-low attachment culture ware (Corning CLS3471) and kept within a humidified incubator at 37°C and 5% CO2. When required, PDN models were dissociated in 1× Accutase (STEMCELL Technologies) for 5 minutes at 37°C.

### 4.3 Human T cell stimulation and sampling

CD8+ T cells were then isolated using positive selection (STEMCELL Technologies 17853), followed by CD4+ T cell isolation using negative selection (STEMCELL Technologies 17951). CD8+ and CD4+ T cells were cultured separately overnight at 37 °C with 5% CO2 in complete human T cell media at a concentration of 10^6^ cells/mL. The following day, T cells were activated using CD3/CD28 Dynabeads (Thermo Fisher 11131D) at a bead:cell ratio of 1:1. Beads were removed from cells after 48 hours of stimulation. T cells were maintained in culture for 15 days at a concentration of 0.5 x106 5*×*^5^ cells/mL, with their media replenished every 48 hours. At each sampling time point, 10^4^ cells were transferred to a skirted 96 well plate and diluted 1:1 in 2x Buffer TCL for a total volume of 50 *μ*L. Cells were mixed until lysed, pulse spun and stored at -80 °C until required for use. Each time point was sampled three times each from two individual donors.

### 4.4 Murine dendritic cell differentiation

BM cells were enriched for cKit+ fraction using anti-CD117 magnetic beads and MACS MS columns (Miltenyi Biotech 130-042-201). The cKit-enriched BM cells were then cultured in complete RPMI with Flt3L-Ig 0.8 *μ*g/mL (BioXcell BE0098), as 3 biological replicates. After 5 days in culture 5,000 - 10,000 cells from the following subsets were FACS sorted directly into 40 *μ*L 2xTCL buffer with either a FACS Aria III or Aria W (BD Biosciences): CDP (CD11c–MHCII–SiglecH–), pre-cDC (CD11c+MHCII–SiglecH–), pre-pDC-like (CD11c+MHCII–SiglecH+CCR9–) and cDC2 (Siglec-H–CD11c+MHC-II+XCR1–Sirp*α*+). Similarly on day 7, cultures were additionally sorted for pDC (CD11c+MHC-II+Siglec-H+CCR9+) in addition to the same subsets as day 5. FACS antibodies are listed in (Table S2).

### 4.5 Temozolomide treatment of patient-derived neurospheres

PDNs were dissociated into single cell and seeded at 1,000 cells per well in 80 *μ*L media across 2 sister 96-well plates. Plates were then observed until individual spheres formed and exponential growth observed. For each plate, 20 *μ*L of 7 serially diluted concentrations of TMZ (Sigma-Aldrich, T2577, 100 *μ*M - 31.6nM) or vehicle control were then added to each well. This 20 *μ*L media also contained a 1:1000 dilution of the Caspase 3/7 Red dye (Invitrogen, C10430). Following this, one plate was placed within the IncuCyte live cell imaging microscope, and the other within a separate incubator to be used for TIRE-Seq. Within the IncuCyte, images were taken every 4 hours for 7 days. For the TIRE-Seq plate, samples were lysed with 2x Qiagen buffer TCL at 3, 5, and 7 days post-treatment and stored at -80°C until processing.

### 4.6 Modified Prime-seq

Prime-Seq was largely conducted as per Janjic et al., [14] with minor changes listed in Supplementary Method C.5. The complete protocol is available at protocols.io.

### 4.7 TIRE-seq

Full details of the protocol are given in Supplementary Method C.6 and protocols.io. Briefly, cells were lysed in TCL buffer (Qiagen 1070498) and mRNA was captured with a Turbocapture plate (Qiagen 72251). Reverse transcription and linear amplification were conducted directly on immobilised mRNA. Soluble cDNA is then pooled, harvested and concentrated prior to a semi-random hexamer extension reaction to introduce Illumina sequencing primers. Two rounds of PCR are then conducted to complete Illumina based library preparation.

### 4.8 Sequencing

Short-read sequencing was performed on the Illumina NextSeq 2000 instrument on a P2 flow cell. The sequencing configuration for the modified implementation of Prime-seq was Read 1: 28nt, Index 1: 8nt, Index 2: 8nt, and Read 2: 90nt with 1% PhiX DNA. The sequencing configuration for TIRE-seq experiments were Read 1: 50nt, Index 1: 8nt, Index 2: 8nt and Read 2: 72nt with 1% PhiX DNA.

Long-read sequencing was conducted by ligating full length Prime-seq cDNA with nanopore adapters (Oxford Nanopore Technologies SQK-LSK114). The sample was loaded on a PromethION Flow Cell (R10.4.1) and sequenced for 3 days.

### 4.9 Bioinformatics preprocessing

Sequencing data was processed into count matrices with zUMIs v2.9.7 version [38]. Reads were mapped against 10x Genomics pre-built references (July 7, 2020 version); GRCh38 v2-7-3a for human experiments, mm10-2020-A for mouse experiments or hg19 and mm10 for mixed species experiments. Long-read data was preprocessed using FLAMES v2.0.1 [39].

### 4.10 Bioinformatics analysis

All downstream analysis was performed in R version 4.4.1 [40]. The code, data and analyses used to generate these figures is available from GitHub. Tabular data was manipulated with the tidyverse package [41].

### 4.11 Statistical analysis

Differential gene expression analysis was performed with edgeR version 4.2.1 with established workflows based on case-control or time course experimental designs [42, 43].

### 4.12 Data and code availability

The count matrices and code for this manuscript are available from GitHub. Raw fastq data will be made available upon publication through a data access committee.

Prime-seq data for benchmarking was downloaded from Array Express (E-MTAB-10142) [14].

## Supplementary information

Supplementary information includes a detailed methods section, a supplementary note, supplementary table 1-2 and supplementary figures 1-4.

## Acknowledgments

We thank all those individuals who donated the samples that enabled this study. We acknowledge the WEHI Advanced Genomics Facility, Flow Cytometry Facility and Animal Bioservices for professional and timely service. We thank Joan Heath for feedback on our manuscript. D.V.B is supported by funding to the Advanced Genomics Facility from the Walter and Eliza Hall Institute. S.A.B is supported by a VCA mid-career research fellowship (MCRF22003). Z.M is supported by a Greg Lange Fellowship. M.R.J is supported by a NHMRC Investigator Grant (APP2032982). Y.N is supported by a Malaghan Institute Fellowship. Protocol development was funded by the WEHI New Medicines and Advanced Technologies (NMAT) theme. This work was financially supported in part through the authors membership of the Brain Cancer Centre and support from Carrie’s Beanies 4 Brain Cancer. Experimental design figures were created using Biorender.com.

## Declarations

### Author contribution

Conceptualization, D.V.B, R.B. Methodology, D.V.B. Data Acquisition, P.O, Y.N, H.S, Z.M, T.M.B, S.W.O, J.J, D.V.B. Data Analysis, D.V.B, D.M.S, C.W, Y.Y, P.F.H. Formal Analysis, D.V.B. Investigation, D.V.B. Writing, D.V.B. Funding Acquisition, D.V.B, R.B. Supervision, D.V.B, R.B, D.AZ, P.H, S.L, M.E.R, R.S.C, M.R.J, C.A, S.H.N, S.A.B, S.F, J.R.W.

### Competing interests

JRW reports research funding from AnHeart Therapeutics to institute; received consulting fees from AnHeart Therapeutics and Servier; being on advisory boards for Roche and Merck; is a data safety monitoring member for Telix Pharmaceuticals. The remaining authors declare no conflict of interest.

### Declaration of generative AI and AI-assisted technologies in the writing process

During the preparation of this work the author used ChatGPT4 and Claude.ai in order to copy-edit. After using these tools, D.V.B reviewed and edited the content as needed and takes full responsibility for the content of the published article.

## Appendix A Supplementary Tables

### A.1 Supp Table 1

List of primer and well barcode sequences used in this study. See excel sheet.

**Table S2.** Antibodies for FACS phenotyping Murine Dendritic cell cultures.

| Antibody | Fluorochrome | Dilution | Supplier | Clone | Cat |
| --- | --- | --- | --- | --- | --- |
| CCR9 | PECy7 | 1/1000 | eBioscience | eBioCW1.2 | 25-1991-82 |
| CD11c | APC | 1/400 | BD | HL3 | 550261 |
| Clec9a | BV421 | 1/100 | BD | 10B4 | 564271 |
| MHC-II | BV650 | 1/1000 | BD | M5/114.15.2 | 563415 |
| Siglec-H | PE | 1/1000 | Biologend | 551 | 129606 |
| Sirpα | PerCPeF710 | 1/400 | eBiosciences | P84 | 46-1721-82 |
| XCR1 | APC Cy7 | 1/400 | Miltenyi Biotech | REA707 | 130-111-375 |

## Appendix B Supplementary Figures

**Fig. S1.**
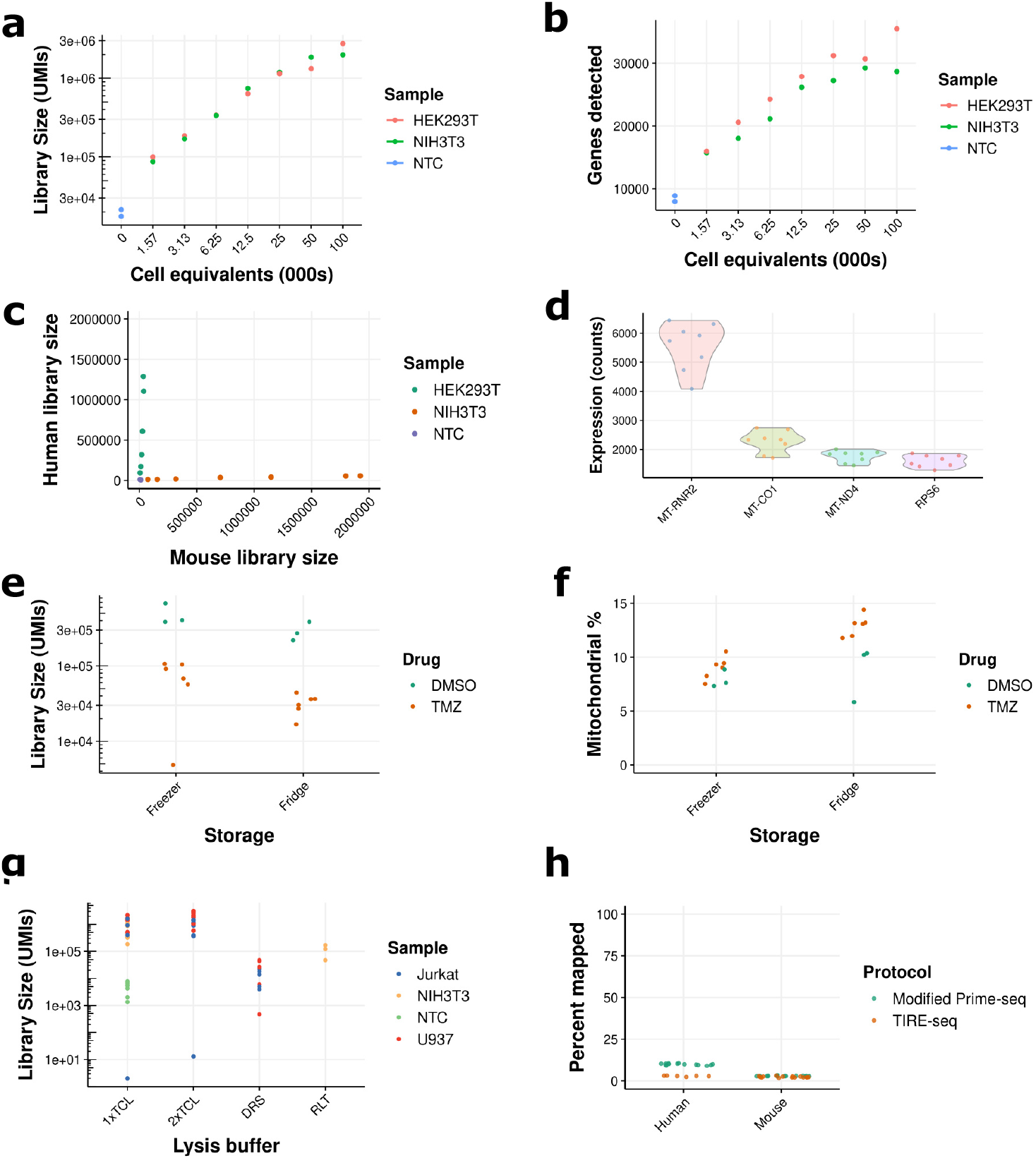
Additional Technical Metrics on the Performance of TIRE-seq. (a) Number of detected molecules (UMIs) across a titration of cell inputs for TIRE-seq. (b) Number of detected genes across a titration of cell inputs for TIRE-seq. (c) Barnyard plot across titrated cell inputs. (d) Highly expressed genes in the TIRE-seq UHRR experiment. Maximum unique combinations with a UMI length of 10 = 1,048,576. (e) Library size comparison in an experiment testing different storage conditions after cell lysis. BT504 cells were treated with 100 *μ*M TMZ for 7 days. (f) Mitochondrial genes expression percentage in an experiment testing different storage conditions after cell lysis. (g) Library size comparison in an experiment testing different lysis buffers on cell lines. 1x, 2x TCL, and RLT = Qiagen buffers; DRS = Zymo DNA/RNA Shield. (h) Mapping rate of sample (header) to the reference genome (x-axis) in Prime-seq compared to TIRE-seq.

**Fig. S2.**
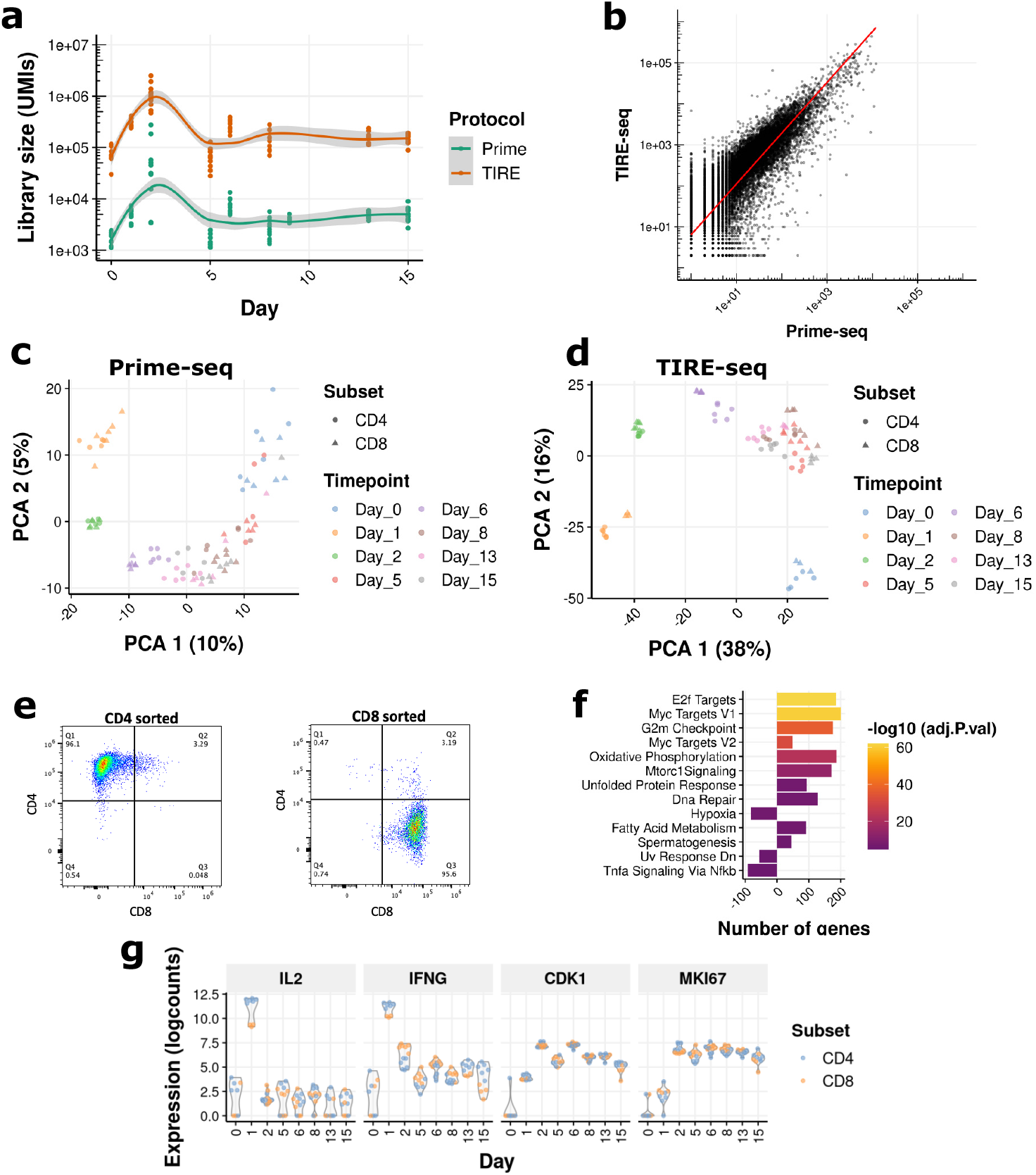
Additional Analysis for the Human T Cell Stimulation Experiment. (a) Number of UMIs detected per timepoint for Prime-seq and TIRE-seq. Reads were downsampled to the same value. (b) Gene expression correlation between Prime-seq and TIRE-seq. (c) Principal component analysis (PCA) of the Prime-seq data for the human T cell experiment. (d) PCA analysis of the TIRE-seq data for the same samples as c. (e) Representative flow cytometry sorting gates for human T Cell Stimulation Experiment. (f) Gene set enrichment analysis of hallmark gene sets comparing day 2 versus day 0 T cells following stimulation. (g) Expression profiles of selected genes across experimental days.

**Fig. S3.**
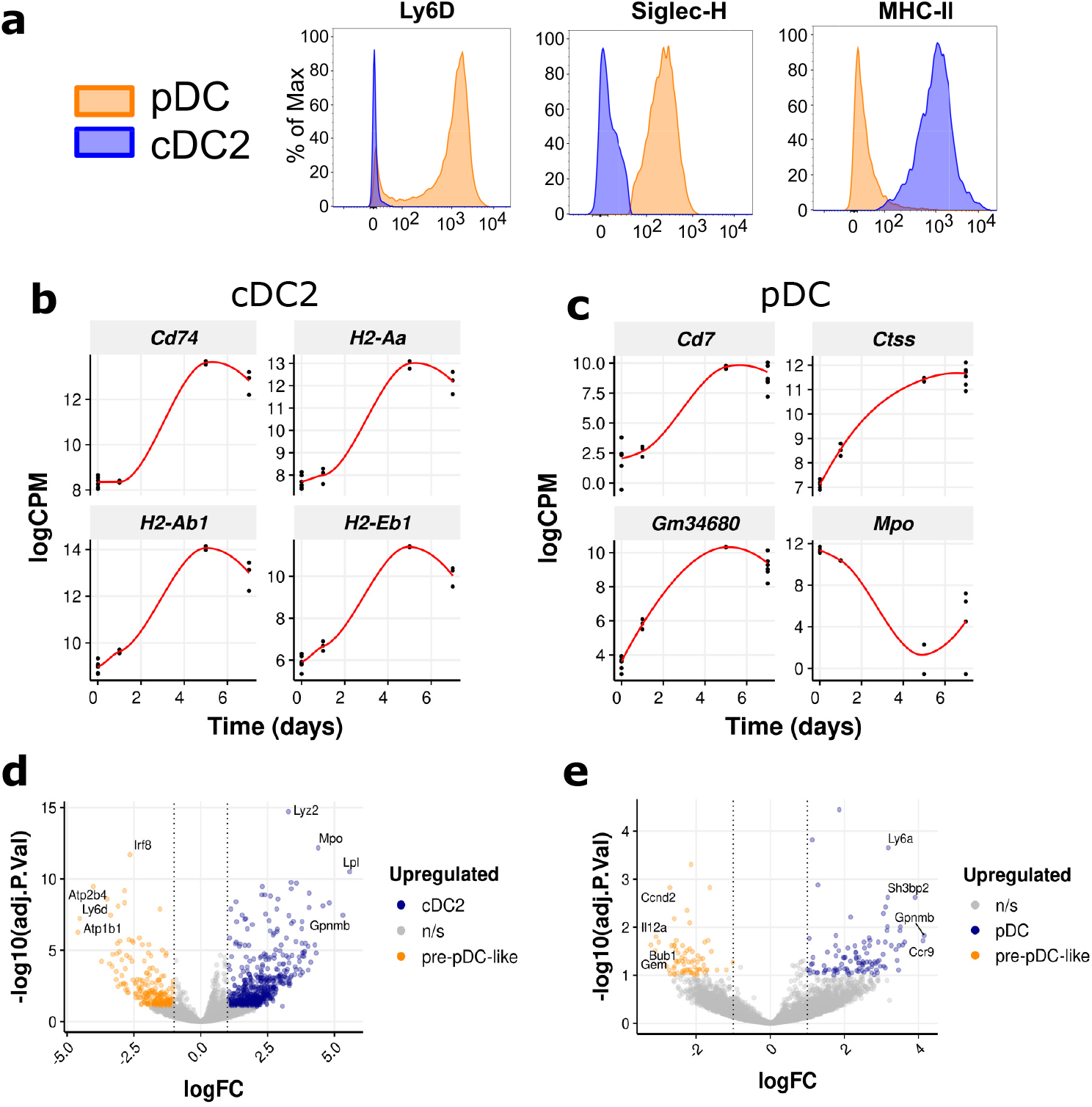
Additional Analysis for the Mouse Dendritic Cell Stimulation Experiment. (a) Flow cytometry data for pDCs and cDC2s. Siglec-H and MHC-II were used as markers for sorting. Ly6D was not used for sorting. (b) Most variable genes over time in cDC2 cells. (c) Most variable genes over time in pDC cells. (d) Volcano plot of differentially expressed genes between cDC2 and pre-pDC-like cells. (e) Volcano plot of differentially expressed genes between pDCs and pre-pDC-like cells.

**Fig. S4.**
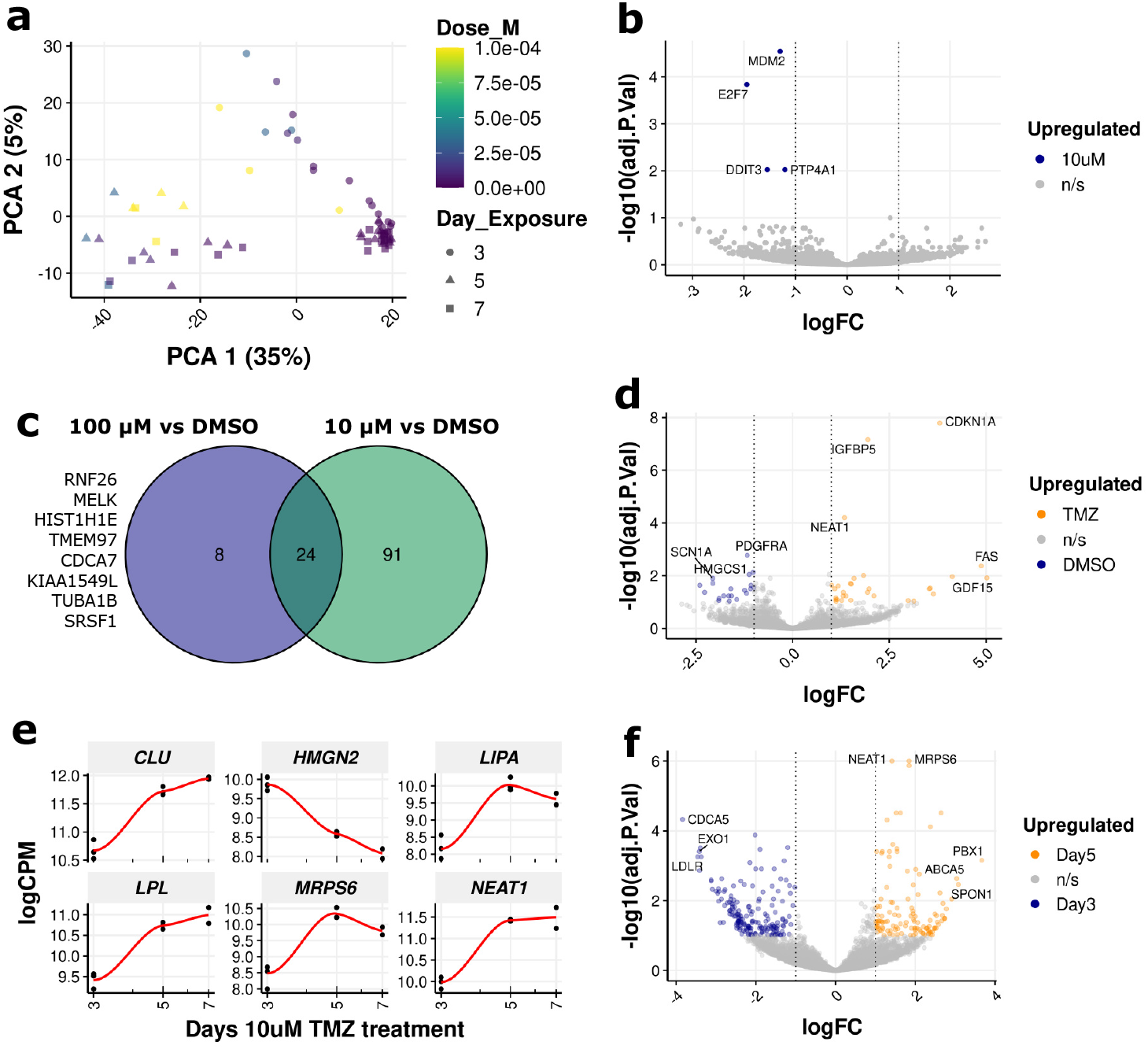
Additional Analysis for TIRE-seq Experiments with Patient-derived Neurospheres. (a) Principal component analysis of TMZ-treated PDNs. Colors represent TMZ molar concentrations. (b) Volcano plot of differentially expressed genes between 100 *μ*M and 10 *μ*M TMZ following 3 days of treatment. No genes were upregulated with 100 *μ*M TMZ treatment. (c) Venn diagram comparing differential expression results between the indicated TMZ concentrations. Unique differentially expressed genes between 100 *μ*M TMZ and DMSO are listed, all of which are downregulated (upregulated in DMSO). (d) Volcano plot of differentially expressed genes between 100 *μ*M TMZ and DMSO following 3 days of treatment. (e) LogCPM values of the top six most variable genes over time with 10 *μ*M TMZ treatment. (f) Volcano plot of differentially expressed genes between day 5 and day 3 timepoints treated with 10 *μ*M TMZ.

## Appendix C Supplementary Methods

### C.1 UHRR Benchmarking Experiment

The experimental design outlined by Janjic et al. was followed with modified volumes [14]. Stock solutions consisted of 100 ng/*μ*L Universal Human Reference RNA (ThermoFisher QS0639, lot 400888-000) and a 1:1000 dilution of ERCC Mix 1 (ThermoFisher 4456740). For both the modified Prime-seq and TIRE-seq protocols, a master mix sufficient for eight replicates was prepared and subsequently aliquoted onto the respective input microwell plates. In the modified Prime-seq protocol, each reaction volume contained 11 ng of UHRR and a 1:10,000 dilution of ERCC Mix A in 1.25 *μ*L of water. In the TIRE-seq protocol, each reaction volume contained 11 ng of UHRR and a 1:10,000 dilution of ERCC Mix A in 20 *μ*L of 1x Qiagen TCL buffer.

### C.2 TIRE-seq lysis buffer comparison

For the 1x TCL and RLT conditions, 250,000 cells were pelleted by centrifugation at 300 g for 5 minutes at 4 °C. The supernatant was removed, and the pellet was washed twice with 1 mL of PBS. The pellet was then resuspended in 500 *μ*L of 1x TCL (Qiagen 1031576) or RLT Plus (Qiagen 1053393). For the 2x TCL or DNA/RNA Shield conditions, 1 mL of 2x TCL (Qiagen 1070498) or 1 mL of 2x DNA/RNA Shield (Zymo Research R1200-25) was directly added to 500,000 U937 or Jurkat suspension cells in 1 mL of RPMI + 10% FCS media. Lysates were mixed thoroughly before storage at -80 °C. For the TIRE-seq experiment, 20 *μ*L of lysate (10,000 cell equivalents) were processed.

### C.3 PBMC and T cell isolation

PBMCs isolated by Ficoll gradient separation. Isolated PBMCs were then cryopreserved in FCS with 10% DMSO at a concentration of 5 *×* 10^7^ cells/mL. Human healthy donor PBMCs were thawed and rested overnight at 37 °C in complete human T cell media (RPMI-1640 (Gibco) supplemented with 10% FCS (Bovogen), 1 mmol/L sodium pyruvate (Gibco), 2 mM GlutaMAX-I (Gibco), 0.1 mM non-essential amino acids (Gibco), 50 *μ*M Beta-mercaptoethanol (Sigma) penicillin-streptomycin (Gibco) and 50 IU/mL rhIL-2 (Peprotech, 200-02)) at a concentration of 10^6^ cells/mL.

### C.4 RNA extraction

For small-scale Prime-seq experiments (Fig. 1), cells were lysed in Qiagen Buffer RLT and RNA was extracted using the Qiagen RNeasy Plus Mini Kit (Qiagen 74134), including the gDNA Eliminator column, following the manufacturer’s instructions. RNA was quantified with a Qubit RNA High Sensitivity Assay (Thermo Fisher Q32852), and the RNA Integrity Number (RIN) was evaluated using a TapeStation RNA High Sensitivity Assay (Agilent 5067-5579).

For the high-throughput Prime-seq experiment (Fig. 2), 10 *μ*L of cell lysate was transferred to a 384-well V-bottom plate (Eppendorf 0030623304), followed by the addition of 10 *μ*L of water using a BenchSmart semi-automated dispenser (Eppendorf). 36 *μ*L of SPRI beads (1.8x ratio) were added to each well, mixed, and incubated for 5 minutes at room temperature. The plate was placed on a magnet for 5 minutes at room temperature, and a BlueWasher (BlueCatBio) was used to remove supernatants and perform three washes with 50 *μ*L 80% ethanol. After 2 minutes of drying at room temperature, RNA was eluted in 8 *μ*L of water.

A FlexDrop (Revvity) was used to dispense 1 *μ*L of DNase I 10x buffer, followed by 1 *μ*L of DNase I enzyme. Samples were mixed by vortexing and incubated at 37 °C for 15 minutes. 1 *μ*L of 0.5 M EDTA was next dispensed using the FlexDrop, followed by incubation at 75 °C for 10 minutes and cooling on ice. SPRI beads were resuspended by vortexing and 11 *μ*L of 100% isopropanol was added to each well. Samples were incubated for 5 minutes at room temperature, followed by another 5 minutes on a magnet. The supernatant was removed by BlueWasher and three washes with 50 *μ*L of 80% ethanol were performed, followed by 2 minutes of drying at room temperature. RNA was finally eluted in 8 *μ*L of water. A 1.2 *μ*L aliquot of this RNA was used for the Prime-seq experiment.

### C.5 Modified Prime-seq implementation

Prime-seq was largely conducted as described by Janjic et al. [14], with the following modifications:

The reverse transcription primer omitted the SMARTer sequence and followed the structure: TruSeq read 1 - 18 nt UMI – 10 nt well barcode. UMIs incorporated the YR YR structure from Karst et al., to enhance sequencing accuracy on the Oxford Nanopore platform [44]. Well barcodes were designed with a minimum Levenshtein distance of 4. Sequences were filtered by Primer3 for Tm within 2 degrees of each other and ΔG *>* -1500 cal/mol for both hairpin and homodimer structures [45]. All reverse transcription primer sequences are provided in Table S1.

At the protocol level, experiments were conducted in 384-well plates (Bio-Rad HSP9901), and volumes were reduced to one-quarter of the original. Library preparation followed the protocol by Janjic et al., with the exception of using the *tenX adaptTop* and *tenX adaptBot* adapters during the ligation step (Table S1). This modification enabled the use of the Dual Index Kit TT (10x Genomics 1000215) in the library PCR step, utilizing existing surplus reagents in our laboratory. The detailed protocol is available at protocols.io.

### C.6 TIRE-seq

A step-by-step protocol is available at protocols.io. All primer sequences are available in Table S1. 2x Qiagen buffer TCL (Qiagen 1070498) was added at a 1:1 ratio with culture media in a total volume of less than 80 *μ*L before storage at -80 °C. A BenchSmart 96 semi-automated pipet head was used to mix and transfer 20 *μ*L of lysed cell samples to a TurboCapture 96-well mRNA Plate (Qiagen 72251). RNA was hybridized to the TurboCapture plate at room temperature on an orbital shaker for 60 minutes, after which all liquid was removed.

A 20 *μ*L reverse transcription mixture, consisting of 1x Maxima RT buffer, 1 mM dNTPs, 1 *μ*M Turbo STD TSO, 0.2 U/*μ*L SUPERaseIN, and 100 U Maxima H-Reverse Transcriptase, was added to the TurboCapture plate. Reverse transcription was performed in a thermocycler with the following program: 25 °C for 15 minutes, 50 °C for 45 minutes, and 80 °C for 15 minutes. Following incubation, 150 *μ*L of 0.1 M NaOH was added to the mixture for inactivation. To remove mRNA and expose first-strand DNA, all liquid was removed, followed by the addition of 100 *μ*L of 0.1 M NaOH for 5 minutes at room temperature followed by two washes with 10 mM Tris pH 7.5. Immobilized first-strand cDNA was converted into soluble second-strand cDNA via linear amplification with 1x KAPA HiFi and 0.5 *μ*M Turbo SSS Well Barcoding Primer in a final volume of 20 *μ*L. Linear amplification was performed with the following program: initial denaturation at 98 °C for 2 minutes, followed by 10 cycles of denaturation at 98 °C for 20 seconds, annealing at 60 °C for 30 seconds, and extension at 72 °C for 30 seconds, with a final elongation at 72 °C for 2 minutes.

Samples were pooled and concentrated using SPRIselect beads (Beckman Coulter B23318), diluted 1:4 in SPRI buffer (10 mM Tris base, 1 mM EDTA, 2.5 M NaCl, 20% PEG 8000, 0.05% Tween 20, pH 8.0) at a 1:1.2x ratio, and eluted in 20 *μ*L of water.

To reduce the length of ssDNA for short-read sequencing, a Klenow reaction was performed with 1 *μ*M semi-random TURBO rand v2 strand oligo in a final volume of 20 *μ*L. The first PCR amplification was carried out with 400 nM Turbo v2 PCR1 Fwd and Rev primers in a 50 *μ*L volume. The cycling conditions were: initial denaturation at 98 °C for 2 minutes, followed by 10 cycles of denaturation at 98 °C for 20 seconds, annealing at 65 °C for 30 seconds, and extension at 72 °C for 30 seconds, with a final elongation at 72 °C for 2 minutes. The first PCR product was purified with a 1x SPRI bead ratio, and one-quarter of the product was used as input for a second PCR reaction.

The second PCR amplification was performed in a 25 *μ*L volume with 5 *μ*L of SI-TT index primers (10x Genomics PN-1000215). The cycling conditions were: initial denaturation at 98 °C for 2 minutes, followed by 8 cycles of denaturation at 98 °C for 20 seconds, annealing at 54 °C for 30 seconds, and extension at 72 °C for 20 seconds, with a final elongation at 72 °C for 2 minutes. The PCR2 product was purified using a 0.8x SPRI bead ratio.

## Appendix D Supplementary Note

### D.1 Reproducing the UHRR benchmarking experiment

Our modified implementation of Prime-seq exhibited lower performance compared to the dataset reported by Janjic et al., (E-MTAB-10142) (Fig. 1c) [14]. While we made extensive efforts to reproduce their experimental conditions and adhered closely to the dilution protocol, we note that certain aspects of the protocol were not fully documented. Our Universal Human Reference RNA (UHRR) reagent was freshly procured within one month prior to the experiment, whereas our ERCC stock was over five years old. Given the high cost of repurchasing ERCC stock for a single experiment, we opted to use the older stock, which may have contributed to discrepancies in performance.

One notable difference in our modified Prime-seq protocol was a reduction in reaction volumes by one-quarter, coupled with the use of 384-well plates instead of 96-well plates. Additionally, in our Prime-seq reverse transcription primer design, we omitted the SMARTer sequence at the 5’ end of the primer to significantly reduce oligo synthesis costs by shortening the primer length below 90 nucleotides. The SMARTer sequence is designed to enable semi-suppressive PCR during the first full-length cDNA amplification, thereby minimizing the formation of short amplicons.

For the cell lysate experiment (Fig. 1d), we note that our experiments were conducted using a different stock of HEK293T cells than Janjic et al., Cell line authentication was performed via STR typing, and we confirmed the absence of mycoplasma contamination using FastQ Screen. We believe that comparing TIRE-seq with our modified implementation of Prime-seq is the most appropriate approach, as both datasets utilized identical lysates, ensuring consistency in the biological material.

Given the major innovation of TIRE-seq is the integration of RNA extraction with library preparation, we do not think purified RNA is a worthwhile input to this protocol. TIRE-seq is best suited to high throughput cell culture experiments in microwell plates where lysis can be achieved by the the addition of an equal volume of Qiagen 2x TCL buffer to cell culture media.

## Notes

https://github.com/WEHIGenomicsRnD/Manuscript_TIRE_seq

https://dx.doi.org/10.17504/protocols.io.j8nlk8rqdl5r/v1

https://dx.doi.org/10.17504/protocols.io.kxygx34qog8j/v1

